# Real-time feedback control microscopy for automation of optogenetic targeting

**DOI:** 10.1101/2025.08.17.670729

**Authors:** Lucien Hinderling, Alex E. Landolt, Benjamin Grädel, Laurent Dubied, Cédric Zahni, Moritz Kwasny, Dario Bassi, Agne Frismantiene, Talley Lambert, Maciej Dobrzyński, Olivier Pertz

**Affiliations:** Institute of Cell Biology, University of Bern, Baltzerstrasse 4, 3012 Bern, Switzerland; Graduate School for Cellular and Biomedical Sciences, University of Bern, Switzerland; Departments of Systems Biology and Cell Biology, Harvard Medical School, Boston, MA, USA

**Keywords:** smart microscopy, microscope automation, closed-loop control, optogenetics

## Abstract

Optogenetics has revolutionized our ability to study cellular signaling by enabling the precise control of cellular functions with light. Most classical implementations rely on fixed or manually updated illumination patterns, limiting their ability to accommodate living systems that move, change shape or rapidly adapt their signaling states. Here, we present an experimental platform for Feedback Adaptive Real-time Optogenetics (FARO) that combines automated image segmentation, tracking, feature extraction, and adaptive hardware control to dynamically adjust optogenetic stimulation based on live cell behavior. By continuously analyzing biosensor signals, FARO updates illumination patterns in real time across biological scales; from maintaining stimulation on specific subcellular regions, to selectively activating single cells in deforming tissue. This fully automated, Python-based framework is built on open standards for data management and microscope control and data handling, supporting large-scale experiments and long-term timelapse studies, compatible with different microscope hardware. Eliminating the need for human intervention to reposition light patterns or select target cells enables reproducible, systematic and high-throughput interrogation of spatiotemporal signaling. We show how automated and adaptive optogenetic perturbations are a powerful tool to study how local signaling events shape cellular behavior, from subcellular dynamics and single-cell migration up to emergent tissue-level processes.

## Introduction

Cells are dynamic systems, constantly evaluating and responding to their environment through tightly regulated signaling networks. Traditional representations of signaling pathways oversimplify these processes, ignoring spatial organization, stochastic fluctuations, and mechanical coupling between molecules, organelles, and cells. To fully understand cellular behavior, researchers must be able to not only image but also perturb signaling in both space and time. A major challenge in perturbing signaling pathways is cellular adaptation. Long-term genetic and pharmacological perturbations often trigger compensatory mechanisms, making it difficult to isolate direct effects [1, 2]. Feedback loops and crosstalk can restore homeostasis, masking the true role of a given pathway. Optogenetics provides a solution by enabling acute, localized perturbations that do not trigger these compensatory responses. Unlike genetic loss-of-function (knockout, knockdown), or chemical inhibition approaches that do not allow for precise spatio-temporal control, light-sensitive protein switches enable precise, reversible control of biochemical activities within seconds with micrometer resolution [3, 4]. Combining optogenetic actuators with spectrally compatible biosensors creates actuator– biosensor circuits that enable systematic probing of signaling input–output relationships at spatiotemporal scales inaccessible with classical methods that average over cell populations or capture only steady-state snapshots. For example, pairing an optogenetic receptor tyrosine kinase with an ERK biosensor allows to activate growth factor signaling and measure downstream ERK activity dynamics at the single-cell scale [5, 6]; similarly, combining optogenetic guanine nucleotide exchange factors (GEFs) with Rho GTPase biosensors and cytoskeletal markers allows to activate Rho GTPase signaling and measure subcellular changes in activity and cytoskeletal organization [7–9]. Throughout this work, we develop and apply such actuator– biosensor circuits across different signaling pathways.

Cells move, change shape, and exhibit heterogeneous behaviors, making static illumination patterns ineffective. Due to a lack of adequate technologies, optogenetic experiments still mostly rely on manually selecting and updating regions of interest (ROIs) for light stimulation, which prohibits large-scale or long-term dynamic studies.

To address these limitations, new microscope platforms must integrate real-time feedback and automation into optogenetic experiments. Advances in smart microscopy provide a solution by combining live imaging and computer vision for adaptive light stimulation [10–15], see [16, 17] for recent reviews. Deep learning-based cell segmentation enables these systems to track and respond to cellular events in real time, allowing fully automated experiments based on user-defined rules. By conditioning stimulation on real-time cellular readouts, such systems can also probe feedback within signaling networks, revealing how the same input produces different outputs depending on the cell’s current state.

Here, we present *FARO* (Feedback Adaptive Real-time Optogenetics), a smart microscopy platform for automated optogenetic stimulation across biological scales. Built on open source software, our system analyzes live images to extract cellular features, makes instantaneous decisions, and delivers targeted light stimulation without human intervention. By combining optogenetics with real-time automation, our approach enables experiments that were previously not accessible, offering new insights into dynamic cellular processes. We demonstrate applications across different cellular systems and signaling pathways.

## Results

### Overview experimental platform

Our experimental platform integrates real-time image processing with automated optogenetic stimulation to enable closed-loop control of living cells (Fig. 1A). In each iteration, the system acquires microscopy images, processes them using a computer vision pipeline to segment and track cells, extracts quantitative features, and generates a stimulation mask that is projected onto the sample using a digital micromirror device (DMD) for optogenetic activation. This cycle repeats continuously, allowing the system to dynamically adapt stimulation patterns to changes in cell behavior.

**Fig. 1.**
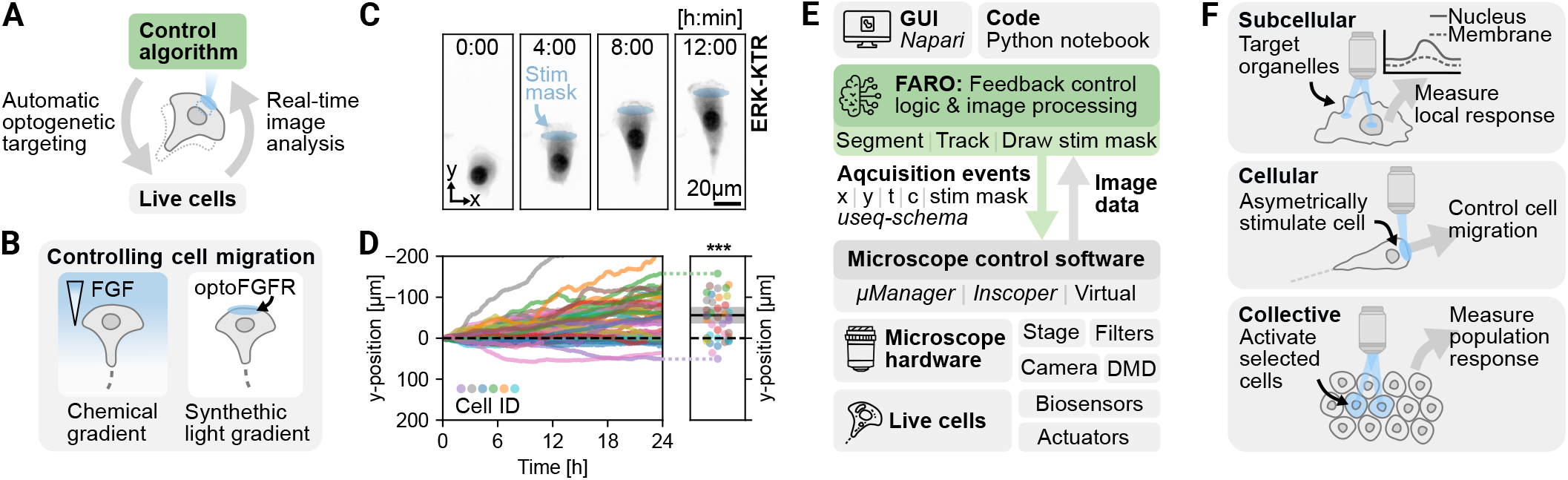
Real-time feedback control microscopy for automated optogenetic targeting. **A:** Closed-loop control of live cells. Computer vision segments and tracks cells in real time, while a control algorithm generates a stimulation mask to dynamically target specific regions of interest with light for optogenetic activation. **B:** As an illustrative example, we induce directed cell migration. Cells orient toward growth factor gradients; here, we simulate such gradients using optogenetic FGF receptor (optoFGFR) with activation restricted to the cell front. As cells move, the stimulation mask must be continuously updated, requiring real-time feedback control. **C:** Example of a single NIH3T3 cell showing polarization and directed migration in response to optoFGFR stimulation. **D:** Cell trajectories normalized to their initial positions. By 24 hours, cells (N = 44) exhibited significant displacement in the y direction (-56.03 ± 68.66 µm, p = 2.60x10^-6^, t = -5.41, df = 43), while displacement in the x direction was not significant (-3.00 ± 36.73 µm, p = 0.59, t = -0.54, df = 43). Violin plot shows mean and 95% CI. **E:** Overview of the experimental platform (software, hardware, live cells). Users interact with the system through a napari-based GUI and Jupyter notebooks. The control algorithm includes a modular image processing pipeline for segmentation and tracking, and logic for optogenetic stimulation, communicating acquisition events to the microscope control software (here pymmcore-plus/µManager, but compatible with any software accepting useq-schema events). The modularity enables the platform to be used with various imaging modalities, cell lines, biosensors, optogenetic actuators, and other photostimulation approaches. **F:** Throughout the paper we demonstrate applications across biological scales: subcellular targeting enables local perturbation of signaling and precise control of cytoskeletal dynamics; single-cell for steering migration; tissue-level manipulation to study collective signaling.

As an illustrative example, we use feedback-controlled optogenetic stimulation to induce directed cell migration in NIH3T3 mouse embryonic fibroblasts (Fig. 1B). Cells interpret extracellular growth factor gradients by generating steeper intracellular signaling gradients, which drive directed migration towards the growth factor source (chemotaxis) [18]; here, we simulate such gradients using an optogenetic actuator for the fibroblast growth factor receptor (optoFGFR) [5] with activation restricted to one side of the cell, inducing a local protrusion and migration in the direction of stimulation. As the cell moves, the stimulation mask must be continuously updated, requiring real-time feedback control (Fig. 1C). By 24 hours, cells exhibited significant displacement in the direction of stimulation, while displacement in the perpendicular direction was not significant (Fig. 1D). While similar previous experiments required manual adjustment of illumination regions [19], using our automated method, we can now automatically control directional migration of hundreds of cells in parallel. This could significantly enhance the throughput and precision of gradient sensing studies. Passmore et al. [14] and Oatman et al. [15] have also recently demonstrated automated migration control, with optogenetic actuators for Rac or EGFR, respectively.

Figure 1E provides an overview of the software, hardware, and biological components of our system. Briefly, users interact with the platform through a graphical user interface and *Jupyter* notebooks. The experiment is controlled through a feedback loop that acquires images at specific stage positions, then processes the images using a modular computer vision pipeline that can be adapted for different experimental needs. Our system interfaces with microscope control software through a hardware-agnostic schema for describing acquisition events, allowing integration with various microscope setups.

The modularity of our approach enables adaptability to different cell lines, optogenetic actuators and biosensors, making it compatible for diverse biological applications. We demonstrate its versatility on multiple scales, from targeting subcellular structures to guiding single-cell migration and controlling collective signaling at the tissue level (Fig. 1F).

### User interaction and experiment setup

Our platform uses Python and Jupyter notebooks to program, test, and run experiments (Fig. 2A). Once a custom workflow is set up, even users with little programming experience can easily modify parameters and swap modular building blocks (microscope hardware configuration, segmentation and tracking methods, stimulation strategies) directly in the notebook. Its interactivity allows users to preview the image processing pipeline and stimulation logic on live data before starting an experiment. The notebook also serves as automatic documentation, ensuring that all parameters are recorded for reproducibility.

**Fig. 2.**
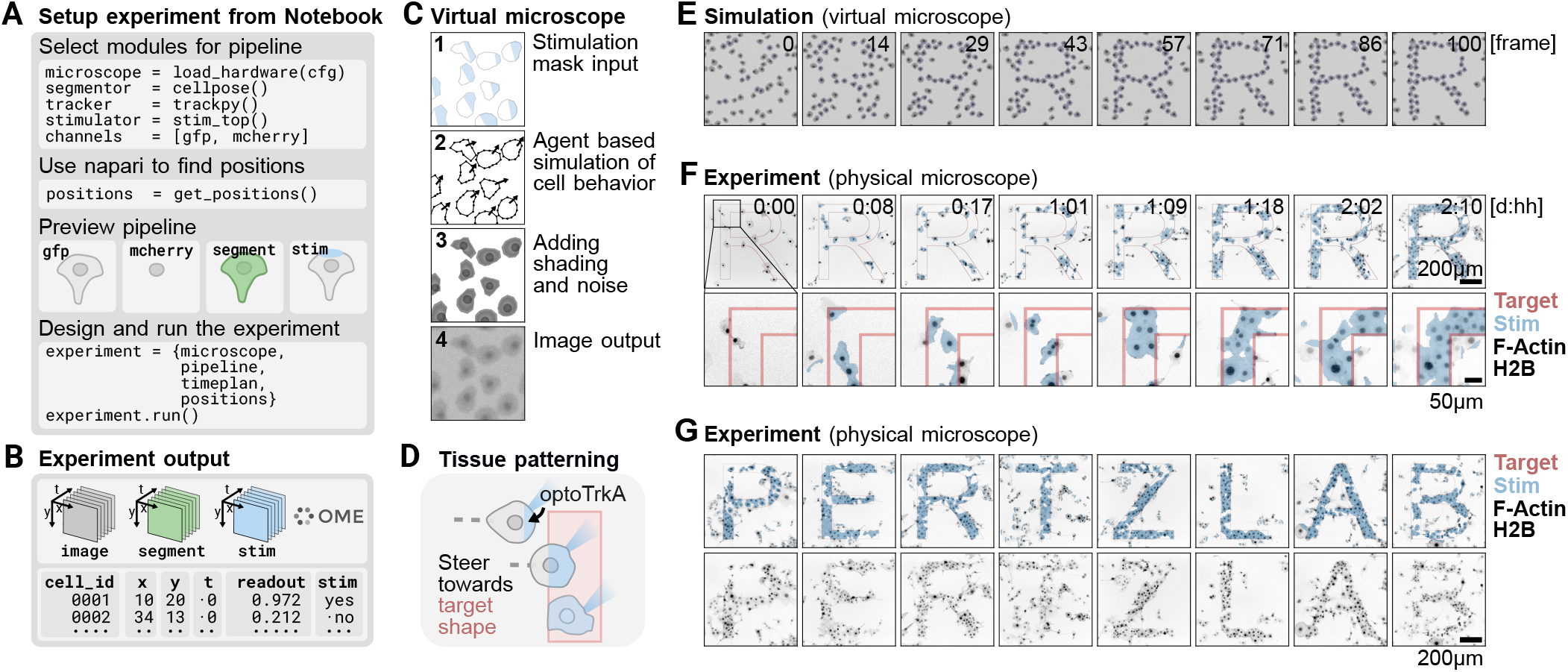
Experiment setup, virtual microscope simulation, and feedback-controlled tissue patterning. **A:** Experiments are set up in Jupyter notebooks with step-by-step instructions to select the relevant modules. The napari interface serves as a graphical user interface to control the microscope, find positions, and directly train the Convpaint classifier on live microscopy data. A preview of the image processing pipeline is displayed. This workflow allows researchers, even with little coding experience, to customize and run experiments. **B:** During the experiment, data is continuously stored, image data and masks in OME-Zarr format, tracking data and other quantitative measurements as pandas dataframes. **C:** Virtual and real microscope hardware can be seamlessly switched to enable pipeline testing and debugging. The virtual microscope includes stage, camera, filters, and spatial light modulator (SLM) components for complete optogenetic experiment simulation. An agent-based simulation generates cell behavior that reacts to spatial optogenetic stimulation. Simulation output is processed with shading and noise to resemble real microscopy data, allowing *in silico* validation of feedback control workflows. **D:** Experimental goal: guide cells that form protrusions in response to optogenetic stimulation towards a user defined shape and maintain them there once they have reached their location. Once the shape is sufficiently occupied, remaining cells in the FOV are steered away from the pattern to increase contrast. **E:** Time series from the simulation, showing cells moving toward the target pattern. 100 frames were simulated in 13 seconds. **F:** Time series from the corresponding real experiment, showing cells gradually forming the same target pattern over several days. Towards the end of the experiment, excess cells are steered away from the pattern. Cells express optogenetic TrkA actuator, biosensors for nucleus and F-actin (H2B / Lifeact). **G:** Example of patterned tissues using the feedback approach; shown are the eight best FOVs and frames manually selected from 24 FOVs of a single experiment. Associated movie (C-G): **Movie S1**.

To control the microscope, adjust focus, and select imaging positions in the sample, we use *napari*, an interactive Python-based multidimensional image viewer [20] with the *napari-micromanager* plugin [21] that provides a user-friendly graphical interface for routine microscope operations like finding sample position or previewing acquisition parameters. Additional napari plugins or custom widgets can be easily integrated, for example, to calibrate the DMD or interactively train image segmentation tools on live data.

Once an experiment is running, data is continuously stored using OME-Zarr [22], along with a data table for additional extracted data (Fig. 2B). The data, including segmentation overlays, tracking information, and stimulation masks, can be directly visualized in napari for real-time quality control. This immediate feedback simplifies troubleshooting and reduces iteration time when an experiment fails. Since data is already extracted during the experiment, there is no need for data processing after the experiment has completed, streamlining analysis and decreasing iteration time between experiments.

### Image processing pipeline

Our experiments rely on real-time image processing to generate segmentation masks, track cells, quantify biosensor activity, and draw ROIs for photoactivation. We selected Python for implementing feedback logic and image/data analysis because of its comprehensive libraries for computer vision and signal processing. Each frame is processed in a separate background thread while the main thread continues to acquire new data. A modular framework allows one to easily replace individual components in the processing pipeline. For segmentation, cell types whose appearance is consistent across experiments or well represented by existing pretrained models can be segmented directly with tools such as *StarDist* [23–25] for nuclear markers and *Cellpose* [26–28] for cytosolic markers. For less conventional cell types, when morphology changes between condition, or subcellular structures, we developed *Convpaint* [29], an interactive pixel classifier that users can train in about one minute using a single representative image taken directly from live data in the napari interface before starting an experiment. To accommodate microscope computers with older hardware (which are often tied to specific hardware configurations and cannot be easily upgraded without breaking compatibility), we also integrate the *Imaging Server Kit* [30], which allows segmentation tasks to be offloaded to more powerful remote workstations or servers. It’s Docker-based deployment simplifies setup while isolating dependencies, thereby avoiding package conflicts between projects or analysis modules. Microscopy images are then transferred to a remote web server via a FastAPI web interface, and the resulting segmentation masks are returned to the FARO pipeline. Cell tracking is performed building on the *trackpy* library [31], which allows new image frames to be added incrementally without reprocessing previous frames, but other tracking tools, such as *btrack* [32], are also compatible with our pipeline. We extract quantitative data, including morphodynamic features such as cell shape, migration speed, as well as signaling activity from biosensor images, enabling targeted stimulation based on live cell behavior. For example, we can selectively stimulate subcellular structures, cells with a specific signaling state or those matching a specific morphology. The data extraction process is highly customizable: users can add their own features by providing any Python function that takes as input the image, segmentation mask and/or tracking data, and returns a stimulation mask and table with the extracted information. The biological processes we studied occur on relatively slow timescales, with image acquisition intervals of at least 10 seconds. In all experiments, throughput was limited by hardware constraints (stage speed, filter wheel switching time, and camera exposure time) rather than image processing speed. To illustrate throughput, a 20x objective provided sufficient resolution to simultaneously control approximately 250 sparsely seeded cells across 24 fields of view (FOVs) at two-minute intervals. In high density epithelial monolayers, we successfully tracked 10’491 cells simultaneously (12 FOVs, 2 min resolution). Although rapid response times were not required for our biological experiments, we tested the system’s ability to quickly process images and respond to detected features in a proof-of-concept experiment and achieved response times below 1 second, including image acquisition, basic image processing (Otsu thresholding, watershed segmentation), mask generation, and hardware adjustments. Also in this case the filter wheel changes were the limiting factor, and could easily be improved by using a dichroic mirror compatible with both the stimulation and acquisition wavelengths.

### Hardware-software integration

Smart microscopy solutions developed in academic labs often encounter interoperability issues between different microscope hardware, because microscope programming interfaces differ significantly between vendors or may be closed-source. In this project, we adopted the hardware-agnostic *useq-schema* [33] to describe image acquisition events: Instead of directly specifying hardware commands, useq-schema defines metadata of acquisition events, such as positions, time points, and imaging channels. The microscope control software then translates this metadata into specific hardware instructions. This approach allows one to abstract the experiment logic from the underlying microscope hardware used. Useq-schema integrates effectively with the open-source microscope control software *µManager* [34, 35] through the Python interface *pymmcore-plus* [36]. This provides FARO access to µManager’s extensive hardware driver library, the largest collection available, enabling compatibility with a wide range of existing microscope setups. For example, different cameras or DMDs, such as the Andor Mosaic 3 and Mightex Polygon 1000, can be used interchangeably, without requiring any change in the code. Importantly, this structured metadata approach also decouples our FARO implementation from any specific microscope control software; preliminary experiments demonstrate that the same experimental protocols can run on microscope control software supporting useq-schema other than µManager, such as UC2 [37] and Inscoper imaging software (https://www.inscoper.com).

### Virtual microscope for testing

Because our platform abstracts hardware behind standardized interfaces, physical and simulated devices can also be swapped seamlessly. Microscopy control scripts are often tested with virtual “fake cameras” that replay pre-recorded images. While sufficient for non-feedback pipelines, such a setup cannot react to perturbations, making it impossible to test feedback control. To overcome this limitation, we developed a virtual microscope that simulates a simple cell model responding to light stimulation. Built on the *UniMMCore* framework of pymmcore-plus, the system allows switching between physical and simulated devices by simply loading a different hardware configuration file, while keeping all device APIs identical. Simulated devices can be implemented entirely in Python without requiring writing C++ based µManager device adapters, lowering the barrier for researchers to create custom virtual instruments. This enables users to develop, test, and debug microscopy experiments in a virtual environment, allowing to systematically verify imaging strategies, feedback loops, and analysis workflows *in silico* before carrying out costly real experiments.

As a test case, we simulated how cells would behave if they expressed an optogenetic tool that induces local protrusions and directional motility, mimicking the behavior seen with optoFGFR discussed earlier (Fig. 1C). We use an agent-based cell simulation, where each cell is modeled as a deformable polygon whose vertices fluctuate under Brownian noise, curvature relaxation, and area conservation. When a stimulation mask is provided, vertices overlapping illuminated pixels protrude outward from the cell centroid and the whole cell receives a directed impulse toward the stimulated side. Cells interact through collisions under periodic boundaries. The scene is rendered as shaded polygons and passed through a contrast, blur, noise, and brightness filter to resemble a microscope image (Fig. 2C). While this model is not physically rigorous, it enables a virtual prototyping workflow in which stimulation strategies are iteratively refined without requiring an explicit mechanical model of cell behavior. In simulations, researchers can compress biological timescales by adjusting the simulation time step, increase the frequency of rare events such as cell divisions to test detection logic, and identify edge cases, all before committing to real experiments.

### Feedback-controlled tissue patterning

We used this simulation to develop an experiment where the goal was to control cell motility to assemble cells into a target shape (Fig. 2D, **Movie S1**). Initial attempts on real cells quickly revealed the difficulty of this problem: naïve strategies, such as simply illuminating cells that already overlap the target, produced no discernible pattern, and each failed attempt consumed an entire day of microscope time. Because the design space of possible stimulation rules is large, exhaustive exploration on the microscope alone was impractical. The virtual microscope removed this bottleneck, allowing us to evaluate dozens of strategy variants in a few hours (Fig. S1). We could even leverage large language models (LLM) to translate high-level instructions, such as *“steer cells only toward free regions of the target”*, directly into updated stimulation logic, visually inspect the result within seconds, and iterate. Mistakes in LLM-generated code were immediately visible and carried no risk of damage to hardware or loss of biological samples. In a few hours, we iteratively addressed problems such as cells crowding one region of the target while leaving others empty, or the pattern contrast degrading once fully formed due to overcrowding. In the final implementation, each cell is guided based on how much it overlaps with the target shape: non-overlapping cells are steered toward unoccupied areas, partially overlapping cells toward a locally expanded portion of the target, and sufficiently overlapping cells are only stimulated in the overlapping region. Once overall occupancy crosses a threshold, the rules switch to a maintenance mode that anchors cells on the pattern while pushing excess cells away. This strategy successfully guided the virtual cells to occupy and maintain the target shape (Fig. 2E). We then deployed the final strategy on real cells. Using a 3 min stimulation/acquisition interval, we ran 24 FOVs in parallel and guided NIH3T3 fibroblasts to assemble into letter shapes (Fig. 2F). We used an optogenetic tool that enables blue light-dependent activation of Tropomyosin receptor kinase A signaling (optoTrkA) [4, 38, 39]. The tool is based on CRY2-mediated light-induced clustering of the intracellular domain of TrkA at the plasma membrane. We generated stable NIH3T3 cell lines co-expressing optoTrkA-mCitrine together with fluorescent reporters for ERK activity (ERKKTR-mScarlet3, not used in this experiment) and cell morphology, specifically markers for nuclei (Histone 2B) and F-actin (Lifeact) [40]. H2B and Lifeact were expressed in the same far-red color channel (H2B-miRFP670nano3, Lifeact-miRFP670nano3). The two signals can be separated digitally due to their distinct appearance/subcellular localization, allowing us to track nuclei for ERK-KTR quantification (which requires a nuclear mask) while simultaneously segmenting cell outlines and visualizing F-actin structures via Lifeact, without sacrificing an additional microscope channel. This allows us to activate TrkA signaling with high spatiotemporal precision and simultaneously monitor how individual cells respond to patterned stimulation through changes in ERK signaling dynamics and morphology.

While the simulator converged within seconds, the actual experiment ran for over two days. Figure 2G shows the eight best FOVs from 24 FOVs of a single experiment, demonstrating that the strategy developed *in silico* translates directly to living cells. Feedback-controlled optogenetics can thus function as a “tissue printer”, dynamically organizing cells into defined configurations without fixed physical scaffolds. Traditional micropatterning approaches to constrain cell collectives in controlled shapes [41, 42] rely on adhesive and non-adhesive proteins/polymers deposited on glass [43, 44], which produces static geometries that cannot adjust to cell heterogeneity or variability in seeding, making it difficult to place an exact number of cells within each pattern. Because our approach continuously monitors and corrects individual cell positions, it can compensate for such variability and e.g. ensure a defined number of cells in each collective. Beyond static arrangements, this feedback-control strategy could enable a form of “dynamic micropatterning”, in which target shapes evolve over time to probe the interplay between signaling and morphogenetic movements during development. In the future, this approach could enable many other applications, such as selectively activating leader versus follower cells during collective migration [19], or controlling cell density to study the propagation of mechanical forces through tissues. In the following sections, we show additional applications of the platform across different biological scales and pathways, focusing more on the biological insights enabled by real-time feedback control.

### Subcellular control of Rho GTPase signaling

Rho GTPases are molecular switches that control cytoskeletal organization: canonically, RhoA drives contractility, Rac promotes protrusion, and CDC42 forms filopodia [45, 46]. However, this classical dogma is increasingly challenged by biosensor imaging studies showing, for example, RhoA activity at the leading edge of protrusions [47, 48]. This complexity arises in part because Rho GTPases are regulated by a large number of GEFs and GAPs [49], many with partially redundant functions, and some even are subject to feedback from the very cytoskeletal structures they control [50, 51]. This complex regulation enables tight spatial patterning of Rho GTPase activity, including dynamic signaling waves at the cellular cortex [52], but also makes traditional genetic perturbation approaches difficult to interpret [53]. Optogenetics in combination with live cell imaging provides a powerful method to investigate these intricate dynamics [54, 55]. Here we show how FARO can leverage optogenetic approaches to control cell shape, automatically target cytoskeletal structures to probe cytoskeletal feedback onto Rho GTPase signaling, and produce signaling patterns with spatial dynamics to replicate phenomena such as propagating waves. While the experiments shown until now used 10x or 20x air objectives with minute-scale acquisition intervals across multiple fields of view in parallel, the experiments in this section were performed with a high-NA 60x oil objective, acquiring fields of view sequentially at much higher frame rates (10 s intervals) to match the spatiotemporal scale of subcellular Rho GTPase signaling and constraints of oil immersion imaging.

#### Controlling cell morphodynamics

Using optogenetic actuators for two different Rho GTPases, Rac and RhoA, we can induce cell protrusion or contractility, respectively, in REF52 rat embryonic fibroblasts (**Movie S2**). We reengineered iLID-based optogenetic actuators for both GTPases (optoTIAM for Rac, based on [9]; optoLARG for RhoA, based on [8]; Fig. 3A). For both constructs we use a high-affinity nanoSspB domain for efficient light-dependent recruitment of the respective GEF DH/PH domain (TIAM or LARG) to iLID at the plasma membrane. We fuse iLid to a stargazin anchor which slows diffusion of the actuator for sharper spatial control of activation [56]. A self-cleaving P2A peptide separates the two components, ensuring equimolar expression from a single construct. To visualize signaling and cytoskeletal responses, we expressed cytoskeletal biosensors for active RhoA, referred to as RhoA* (2xrGBD-dTomato) [57], focal adhesions (paxillin-mCherry) [58], or F-actin (Lifeact-miRFP) [40]. To form protrusions with optoTIAM in a controlled direction, cells were continuously segmented with Convpaint and a stimulation mask was computed at every time point, defined by a cutoff line halfway between the centroid and the bounding box edge in the desired direction of protrusion. By updating the mask to account for morphological changes over time, the stimulation region tracked the evolving cell shape throughout the experiment (Fig. 3B,C). Similarly, automated stimulation of optoLARG induced a gradual buildup of RhoA activity and F-actin at the illuminated edge, followed by retraction of the cell boundary, causing cells to systematically contract on one side (Fig. 3D,E). Both experiments used 10 s acquisition and stimulation intervals. These experiments demonstrate that FARO enables automated control of cell morphodynamics at the subcellular level, opening the door to systematic studies of how spatially defined Rho GTPase inputs are integrated into whole-cell responses.

**Fig. 3.**
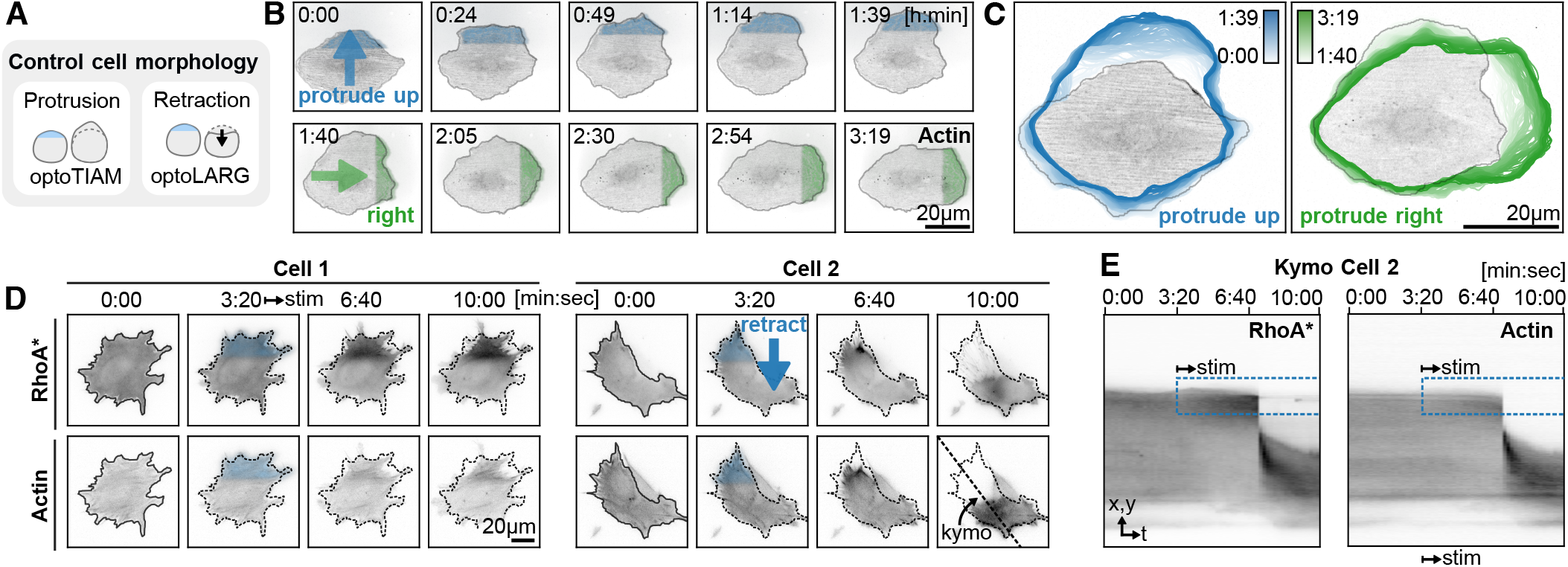
Optogenetic perturbations to control cell shape at the subcellular scale. **A:** Controlling protrusion and retraction using optogenetic actuators for Rac (optoTIAM, panels B,C) and RhoA (optoLARG, panels D,E). **B:** The cell expressing optoTIAM is segmented with Convpaint using an F-actin biosensor image. Light is first projected onto the top of the mask, inducing protrusions in that direction. After 1:39 h:min, stimulation switches to the right side, and the cell redirects protrusions accordingly. **C:** Cell outlines over time show shape deformation. The left panel (0:00 to 1:39 h:min) captures the cell moving upward, while the right panel (1:40 to 3:19 h:min) shows movement to the right. **D:** Cells expressing the optogenetic RhoA actuator (optoLARG) are segmented using Convpaint (dashed outline). The top region is automatically selected for repeated stimulation (projected light shown as a blue overlay on the first stimulation frame), leading to local accumulation of active RhoA (RhoA*) and F-actin, followed by retraction of the stimulated area. Line labeled kymo indicates location of kymograph in E. **E:** Kymograph showing gradual buildup of RhoA* and F-actin at the illuminated edge, then spring-like retraction in Cell 2. Blue box indicates stimulated area. Associated movie: **Movie S2**.

#### Targeting of subcellular structures

Next, we demonstrate automated optogenetic activation of optoLARG with micrometer resolution on specific cytoskeletal structures, here focal adhesions, and measurement of the local RhoA* response (Fig. 4A). A key question in Rho GTPase biology is whether the local cytoskeletal context shapes the response to Rho GTPase activity [50]. To test this, we targeted focal adhesions, known to localize many GEFs and GAPs [50, 59], and paired membrane regions without focal adhesions within the same cell. Cells are segmented using Convpaint on the RhoA* channel and virtually divided into two halves (Fig. 4B): in one half, focal adhesions are detected via the paxillin channel and selectively activated; in the other, non-focal-adhesion regions are selected by sampling near the cell edge with a minimum distance to focal adhesions, providing a paired control within the same cell (Fig. 4C,D). The biological findings are discussed in detail in [58]; here we focus on how FARO enabled the measurements. A critical consideration is perturbation strength: Strong optogenetic activation, as in the previous experiment (Fig. 3D), remodeled the very cytoskeletal structures whose influence on Rho signaling we aimed to measure, making it impossible to disentangle cytoskeletal feedback from the response to the perturbation itself. To preserve the native cytoskeletal context, we used weak, transient stimulations that locally nudged the signaling equilibrium without detectably altering cell mechanics. Although Rho activity appears macroscopically steady, individual molecules undergo continuous cycles of activation and deactivation (signaling flux) [60], and these minimal perturbations allowed us to effectively measure the rates of this flux near steady state. Because such weak perturbations operate in a low signal-to-noise regime, they required the high-throughput automation of FARO, combined with ensemble averaging across thousands of individual adhesions and control regions in hundreds of cells, to achieve statistical significance. With this subtle perturbation regime, we found distinct spatial activity patterns (Fig. 4E), suggesting that focal adhesions provide positive feedback onto RhoA* signaling, consistent with differential regulation by GEFs and GAPs depending on local cytoskeletal context.

**Fig. 4.**
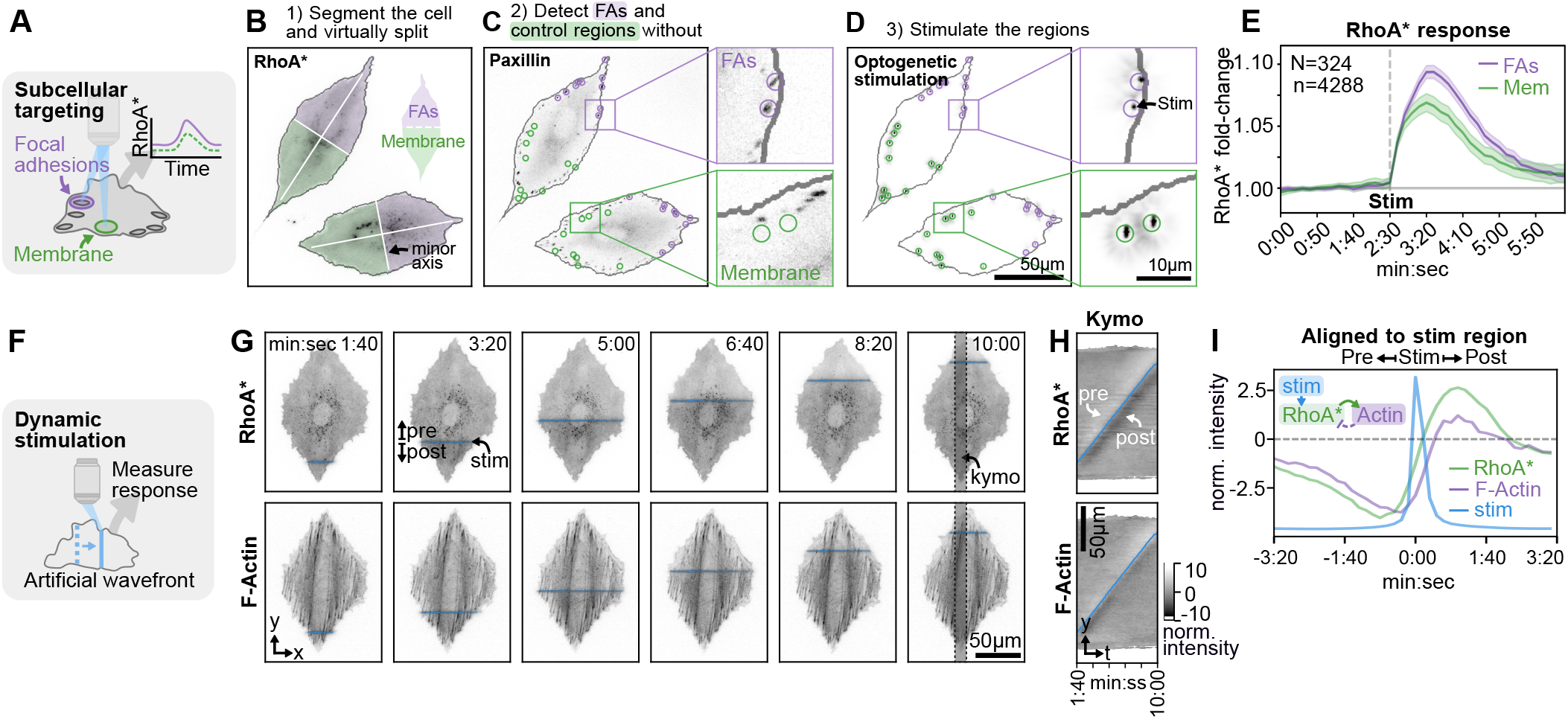
Automatic targeting of subcellular structures and dynamic stimulation patterns. **A:** RhoA is locally activated using an optogenetic GEF (optoLARG) at focal adhesions and membrane regions without focal adhesions. Local RhoA activity (RhoA*) is measured in the stimulated regions using a fluorescent biosensor. Focal adhesions are detected using a paxillin marker. **B:** The cell is segmented using Convpaint on the RhoA* channel and virtually divided along the minor axis. One side is stimulated at focal adhesions, the other at membrane regions, allowing for within-cell controls. **C:** Regions selected for stimulation are marked with circles. **D:** Image of the projected light pattern for optogenetic stimulation. **E:** The process is fully automated, enabling the screening of thousands of stimulation regions (n = 4’288) across hundreds of cells (N = 324). Median and 99% CI are shown. RhoA* shows stronger response at focal adhesions than at the membrane, indicating cytoskeletal feedback on RhoA* signaling. Data is pooled from four technical replicates and was previously reported in [58]. **F:** A line scan induces an artificial wave of signaling activity using optoLARG. RhoA* and F-actin are measured using fluorescent biosensors. **G:** Time series showing RhoA* and F-actin recruitment after stimulation. An image of the projected light pattern is overlaid in blue. Stimulation line width is 1.3 µm (half-height width of the peak-profile). **H:** Kymograph averaged over a 50-pixel-wide region, corrected for photobleaching and structural background, then normalized to mean expression levels. The blue line indicates the y-position of the stimulation mask centroid. RhoA* depletion is observed ahead of the stimulation line, while an increased biosensor signal appears behind it. **I:** Kymograph data aligned to the y-centroid of the stimulation mask, showing dynamics of RhoA* and actin as function of time relative to peak optogenetic activation. The light stimulation profile is manually rescaled and offset for visualization. Scheme shows hypothetical regulatory network that could explain these dynamics. Associated movie (F-I): **Movie S3**.

#### Simulating subcellular signaling waves

Reaction-diffusion systems on the cell cortex generate distinct patterns such as clusters [61] or waves [52] observed in frog oocytes [62], dictyostelium [63], and REF52 cells [64], with the spatio-temporal dynamics of these waves regulating diverse functions. Naturally occurring waves are stochastic, making systematic study difficult. To study their function systematically, we sought to artificially recreate them using optoLARG, replicating the spatial and temporal characteristics of naturally occurring RhoA* waves in a repeatable and controllable manner (Fig. 4F, **Movie S3**). Cells are segmented on the RhoA* channel, and a thin stimulation line is scanned from bottom to top (Fig. 4G). A kymograph analysis, after detrending and normalization, reveals a distinct intensity band trailing the stimulation line for both RhoA* and F-actin, while a band of decreased intensity appears ahead of the wavefront (Fig. 4H). Aligning the measurements to the stimulation region and averaging across time allows a direct comparison of RhoA* and F-actin dynamics. RhoA* precedes F-actin by 30 seconds (three frames) during recruitment, but both reach peak activation simultaneously. After the peak, RhoA* decays at a faster rate than F-actin (Fig. 4I). These dynamics suggest a regulatory loop where optogenetic activation of RhoA* promotes F-actin polymerization, which in turn inhibits RhoA*, possibly through recruitment of a Rho GAP as previously observed [65–67]. Although wavelike optogenetics has been applied at larger scales [68–70], to our knowledge, this is the first demonstration of artificially replicating subcellular signaling waves. Because wave parameters such as propagation speed and wavefront width can be precisely tuned, this approach opens the door to systematic perturbation experiments that could be used to parametrize reaction-diffusion models of cortical Rho GTPase signaling. Combined with the high-throughput automation of FARO, such experiments become feasible at the statistical scale required to constrain these models. As an additional application, the controlled wave signal served as a ground-truth reference to calibrate a computer vision algorithm for automatically detecting spatially coordinated cellular dynamics [64].

### Controlling collective fate decision signaling dynamics in epithelial tissues

Next we show applications of FARO at the tissue level, perturbing the dynamics of MAPK/ERK signaling in 2D epithelial sheets derived from human breast cells (MCF10A). Collective ERK signaling in tissues plays a critical role in wound healing [71–73], morphogenesis [74, 75], and cancer biology [76, 77]. These observations rely on imaging studies that provide single-cell-level activity time series. Optogenetic experiments with static illumination patterns have been applied to whole populations or subgroups of cells, and have shown to be effective in untangling the spatial components of emergent behaviors such as collective movement [78] and feedback regulation within signaling networks [5]. As cells move in tissue, optogenetic control of specific cells requires tracking and adaptive stimulation patterns. We demonstrate that FARO enables phototargeting of single cells during long-term experiments, even using biosensor data extracted in real time to control each cell’s activity based on its signaling history. In the following experiments, we used cells stably expressing a nuclear marker (Histone 2B-miRFP703) for tracking, and an ERK-KTR biosensor (ERKKTR-mRuby2) for measuring single-cell ERK activity based on the ratio of cytosolic to nuclear fluorescence intensity. ERK activity is automatically quantified during the experiment using a multi-step image processing pipeline [79].

#### Inducing targeted cell death

Epithelial tissues maintain integrity through a protective mechanism: when a cell dies, neighboring cells receive a transient ERK pulse that shields them from subsequent apoptosis for several hours, preventing multiple nearby cells from dying simultaneously and forming holes in the epithelium (apoptosis-induced survival [80, 81]). Because spontaneous apoptosis is rare, prior studies often relied on chemotherapeutic agents to trigger cell death, affecting the health of the entire tissue. FARO enables a more targeted approach: by tracking and targeting individual cells with a spot of UV light over several hours (standard 395nm LED in our fluorescence microscope), we were able to induce photo damage leading to cell death in these specific cells (Fig. 5A). The targeted nuclei shrink and are extruded from the epithelium (Fig. 5B), accompanied by a visible contraction of the surrounding tissue (**Movie S4**), triggering a radial ERK activation wave in neighboring cells (Fig. 5C, scheme of how to interpret the ERK-KTR biosensor is shown in Fig. 5D). Because each death event is controlled, its timing and location can be optimized to avoid edge artifacts or overlapping events, increasing the yield of analyzable responses without compromising overall tissue health. While also optogenetic tools for inducing apoptosis exist [82], UV-based photodamage has the advantage of not requiring additional transgenes, and keeping fluorescence channels available for biosensors.

**Fig. 5.**
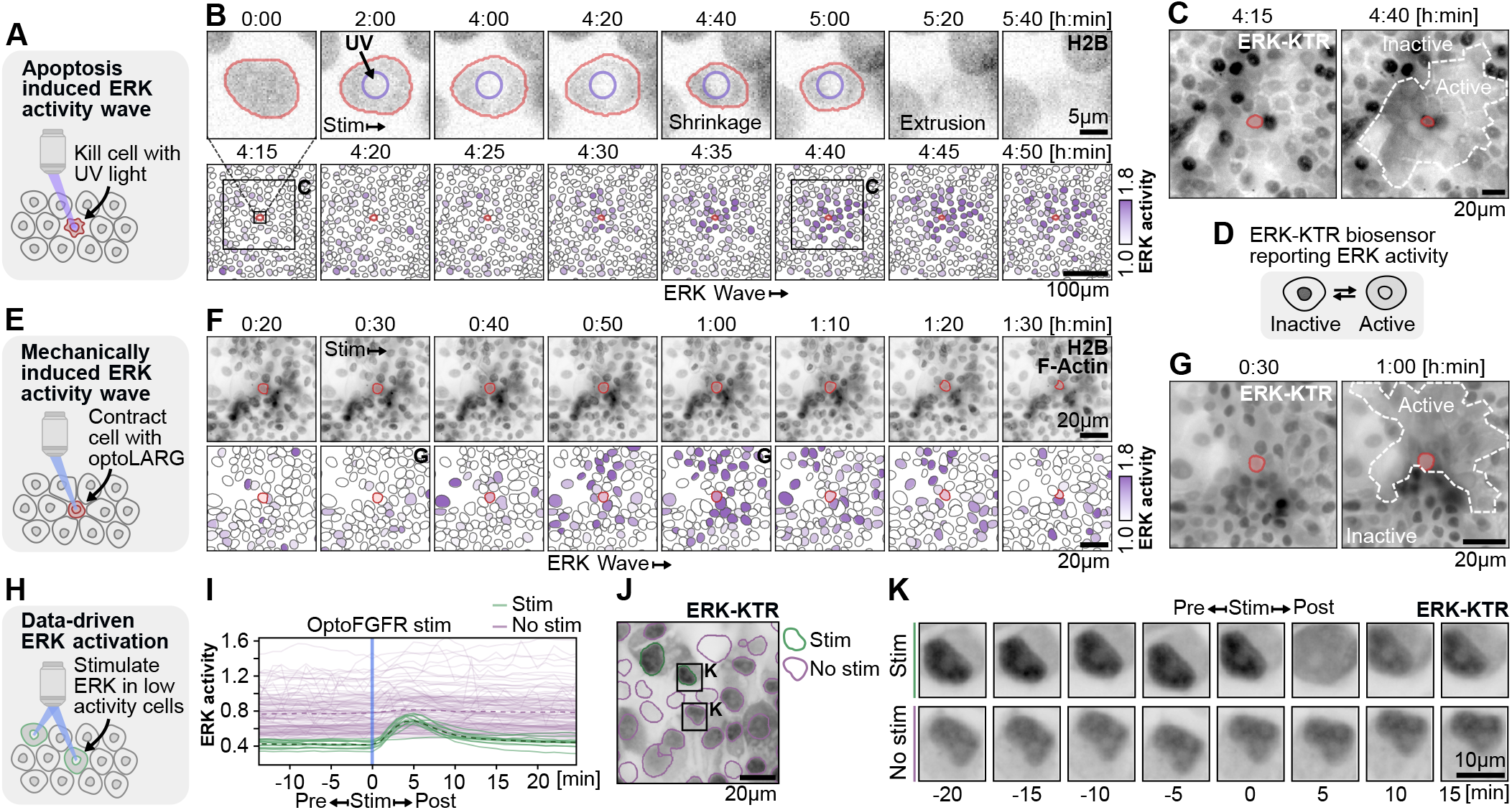
Applications of FARO to study ERK signalling dynamics at the tissue scale. MCF10A cells grown in a confluent monolayer express a nuclear / F-actin marker for tracking (H2B/Lifeact), an ERK activity biosensor (ERK-KTR). **A:** Single cell is tracked and continuously exposed to UV light to investigate how surrounding cells respond to cell death. **B:** Upper panel shows time series centered on the selected cell, showing its mask and the stimulation area (purple circle, ∼5 µm diameter). Bleaching is visible in the stimulated region, followed by nuclear shrinkage and cell extrusion from the epithelium. Lower panel shows zoomed-out view of the epithelium around the time of nuclear shrinkage. Segmentation masks are shown in black, and purple color-code shows ERK activity (ERK-KTR cytosol/nuclear ratio, normalized per cell). The UV-exposed cell is marked in red. The time series shows an ERK activity wave propagating outward from the selected cell. The insets at 4:15 and 4:40 h:min correspond to the crops shown in C. Small inset at 4:15 shows relative size of crops in B. **C:** ERK-KTR signal before (4:15 h:min) and at peak ERK activity (4:40 h:min). The area of active cells is outlined with a dashed line. Active cells are characterized by translocation of the biosensor from the nucleus to the cytosol, as illustrated in D. Note all images are shown with inverted color map (darker is higher camera signal). **D:** Scheme of the biosensor for ERK activity (ERK-KTR). In cells with little active ERK, the sensor is accumulated in the nucleus, and (reversibly) translocates to the cytosol upon ERK activation. **E:** Cells express an optogenetic actuoator for RhoA (optoLARG). A cell with high optoLARG expression is detected and stimulated to contract, mechanically triggering an ERK activity wave in neighboring cells. **F:** Timeseries showing nuclear marker and F-actin stain (top row) and ERK activity (bottom row, normalized per cell). Stimulation starts at 0:30 and is repeated every 20 sec. An ERK activity wave spreads from the selected cell at around 0:50 h:min. **G:** ERK-KTR image at the beginning of stimulation (0:30 h:min) and at peak collective ERK activity (1:00 h:min). **H:** Inducing artificial ERK signaling states in cell collectives expressing an optogenetic actuator for FGFR (optoFGFR). ERK activity is monitored in real time and used to guide optogenetic ERK activation. Cells in the lowest 0.1 quantile of integrated ERK activity are selected for stimulation. **I:** ERK activity (ERK-KTR cytosol/nuclear ratio) plotted over time. Cells with low ERK activity (green) are selected for optogenetic activation. Following stimulation, activated cells exhibit a pulse of ERK activity, whereas unstimulated cells (purple) show only a slight increase (mean activity per group indicated by a dotted line). **J:** ERK-KTR channel image with segmentation outlines indicating whether cells were selected for stimulation. Two cells in close proximity from either class are shown in detail in K. **K:** Time series of two neighboring cells, one selected for stimulation and one not. The stimulated cell shows translocation of the ERK-KTR biosensor to the cytosol after stimulation, while the unstimulated cell exhibits no change in biosensor distribution. Associated movie (A-G): **Movie S4**.

#### Inducing ERK activity wave using optoLARG

Mechanical coupling between contracting apoptotic cells and their neighbors has been observed during ERK wave initiation [80], and ERK is now recognized as a mechanosensitive pathway activated by cell stretch through E-cadherin-dependent ADAM-mediated EGFR ligand shedding [83], notably the same mechanoproteolytic pathway that initiates apoptotic ERK waves [80]. Also ERK waves during wound healing are mechanochemical in nature, with cell stretch and ERK activation propagating in a coupled fashion [69]. Previous optogenetic studies either induced apoptosis directly (OptoBAX) [80, 84] or activated ERK signaling itself to generate synthetic waves [69, 80]. However, whether contractility alone, without apoptotic signaling, is sufficient to trigger an ERK wave has not been causally tested, and the need for optogenetic perturbations to dissect the role of mechanics in ERK wave initiation has been noted [85]. To test this, we mimicked the contraction associated with apoptotic cell extrusion using MCF10A cells expressing opto-LARG (Fig. 5E, **Movie S4**). Using FARO, we automatically identified the cell with the highest optoLARG expression per FOV, expected to generate the strongest contraction upon activation, and stimulated it with a 32 µm diameter spot at 20-second intervals following a 30-minute baseline. This induced rapid contraction of the surrounding tissue toward the targeted cell, followed by an outward-propagating wave of ERK activity (Fig. 5F,G). This demonstrates that mechanical contraction alone, in the absence of caspase activation or release of cellular contents, is sufficient to initiate an ERK activity wave, consistent with mechanical stretch of neighboring cells activating the EGFR ligand shedding pathway that underlies apoptotic ERK waves. This suggests that the mechanical component of cell extrusion directly contributes to triggering the protective survival response.

#### Stimulating cells based on signaling history

FARO can also use single-cell ERK activity data extracted during a live experiment to modulate behavior of each cell based on its signaling history. Epithelial homeostasis and other collective dynamics are emergent behaviors that arise from stochastic single-cell signaling and cell-cell communication [75, 80, 86, 87]. A key open question is whether these collective behaviors can be re-engineered by artificially controlling single-cell signaling dynamics, for example, by equalizing ERK pulse frequencies across a population to test whether signaling heterogeneity is required for tissue patterning. In this proof-of-concept experiment, we measure ERK activity time series and selectively activate cells with the lowest integrated activity using an optogenetic actuator for fibroblast growth factor receptor signaling (optoFGFR) [5, 88] (Fig. 5H). Stimulated cells exhibit a strong ERK activity pulse, while non-selected cells generally show flat activity, with few exceptions (Fig. 5I). These exceptions may result from overlapping protrusions that expose multiple cells to a single illumination spot, or from paracrine signaling between neighbors. Fig. 5J and K show a representative time series of two neighboring cells, one selected for activation and one not. Since cells decode ERK pulse frequency and integrated activity into fate decisions such as proliferation, differentiation, or survival [86, 89–91], adaptive stimulation protocols that impose defined ERK dynamics on specific subpopulations could in principle be used to program fate decisions with single-cell resolution. Extending this approach to larger tissues and additional signaling pathways could enable systematic dissection of the rules governing collective cell behavior.

## Discussion

We present FARO, a real-time feedback control microscopy platform that combines segmentation, tracking, and adaptive light stimulation to enable automated optogenetic experiments across biological scales.

### Experiments enabled by automated feedback control

The applications presented in this work span from subcellular structures to tissue-level collective dynamics. At the single-cell level, inducing directed migration required continuously updating the stimulation mask as cells moved and changed shape (Fig. 1C). Previously, this demanded manual adjustment every few minutes [19], limiting throughput to a handful of cells; FARO simultaneously steered hundreds of cells in parallel (Fig. 1D) and, by tracking each cell’s position against the evolving arrangement of its neighbors, assembled cells collectively into defined tissue patterns over multiple days (Fig. 2F,G). Controlling cell morphology with optogenetic Rac or RhoA actuators relied on real-time segmentation to keep illumination aligned with the evolving cell shape (Fig. 3B-D). At the subcellular scale, probing cytoskeletal feedback on RhoA signaling required automated detection and stimulation of thousands of individual focal adhesions across hundreds of cells (Fig. 4A-E). Artificially replicating subcellular signaling waves required real-time segmentation to precisely position and animate the scanning stimulation line within individual cells (Fig. 4F-I). At the tissue level, inducing targeted cell death required feedback control to keep the UV spot centered on the selected cell as the tissue moved, ensuring only the target cell was killed (Fig. 5A-D). Testing whether mechanical contraction alone can trigger an ERK survival wave required automatically identifying the highest-expressing cell per FOV and maintaining stimulation as the tissue deformed (Fig. 5E-G), disentangling local mechanical from biochemical contributions in a way that bulk perturbations cannot. Selectively stimulating cells based on their real-time signaling history is inherently a closed-loop problem: without continuous biosensor readout and adaptive targeting, such experiments reduce to random or uniform stimulation protocols, which obscures the very heterogeneity they aim to control (Fig. 5H-K). Beyond our own experiments, the platform has been adopted by collaborators for distinct biological questions: Torres-Torres et al. [92] used it to target the front and back of migrating macrophages to investigate signaling specificity among redundant Src family kinases, and Berthoz et al. [93] used it to measure subcellular Rho GTPase dynamics in giant epithelial cells, combining the data with theoretical modeling to study Rac1/RhoA and actin wave interplay. All these examples required continuous segmentation, tracking, and mask generation for every target region at every time point. Whether the challenge is targeting thousands of subcellular structures or maintaining precise stimulation across hundreds of cells over days as the they move and deform, manual intervention quickly becomes impractical. FARO automates this entire loop, enabling experiments at scales and durations that would otherwise not be feasible.

### Technical implementation and interoperability

Commercial solutions for automating microscopy using image feedback currently do not match the requirements of academic research labs, leading to a fragmented ecosystem of custom-built tools that are not interoperable [16]. Here, we use useq-schema to describe image acquisition events: instead of directly specifying hardware commands, FARO outputs useq-schema acquisition events that define the position, time point, imaging channels, and stimulation mask for each frame. The microscope control software then translates these into specific hardware instructions (move stage, change filter, set camera exposure, display DMD pattern), abstracting away vendor-specific implementation details. This means experimental protocols become transferable between labs with different hardware setups, reducing the barrier to adopting published methods and enhancing reproducibility. While all experiments presented here were performed using pymmcore-plus/µManager, preliminary tests demonstrate that our experimental protocols can run on alternative microscope control platforms supporting useq-schema imaging software, without modification to the core experiment logic. For developers, maintaining a single codebase that works across multiple platforms significantly reduces the software maintenance burden compared to supporting platform-specific implementations. Using OME-Zarr as image format further enhances integration with downstream data analysis and storage tools. Limitations remain: the success of useq-schema depends on adoption of the standard, which at the moment remains limited (µManager/Inscoper/UC2), although the wide range of devices supported by µManager means many systems are supported already today. Commercial confocal and super-resolution systems often lack the necessary APIs for external control, restricting this approach mainly to camera-based microscopes. Additionally, the abstraction layer may limit access to vendor-specific advanced features that fall outside the standardized event schema. Despite these challenges, our adoption of useq-schema marks a step toward standardization of smart microscopy approaches. We hope that demonstrating the wide range of applications and benefits of this architecture will encourage broader adoption across both open-source and commercial platforms, contributing to a more interoperable smart-microscopy ecosystem. Beyond interoperability, a practical challenge for any smart microscopy platform is balancing customizability with ease-of-use. We tried to strike a balance by focusing specifically on automating optogenetic targeting, rather than broader smart microscopy applications such as sample tracking or adaptive imaging, and by modularizing components like image segmentation, tracking, and stimulation logic, allowing researchers to tailor the system to their specific research question. A key advantage of our setup is the tight integration with the napari image viewer, providing a graphical interface for experiment setup, real-time visualization, and quality control. While initial setup of an experiment requires coding, for subsequent experiments this integration enables researchers with limited programming experience to run their own experiments by stepping through a Jupyter notebook cell by cell. Markdown cells can serve as inline instructions, guiding users on where to adjust parameters or modify settings. By separating the experiment logic from the data handling code, the experimental logic can also be more easily customized with the help of LLM coding assistants. In the tissue patterning experiment, we used this approach extensively: we described desired changes to the stimulation strategy in natural language, had an LLM generate the corresponding code, and tested variants in the virtual microscope within seconds (Fig. 2E,F). This workflow lowers the entry barrier for advanced customization and dramatically shortens the iteration cycle for developing new experimental strategies. Several smart microscopy platforms for feedback-controlled experiments have been developed in parallel [10–15, 94–102]. To help readers identify which platform best fits their experimental needs and existing hardware, we maintain a comparison and compatibility matrix of both academic and commercial platforms at https://smartmicroscopy.github.io/implementations.html [16], which is community-editable and open for contributions so it can evolve with the field.

### Future directions

While our primary focus is on optogenetic control, the platform is compatible with any light-based perturbation system, e.g. as shown with UV-induced cell death (Fig. 5A,B). We also use the same UV projection setup for microfabrication [103]; similar feedback-control approaches have been applied to maskless lithography [104], highlighting potential uses beyond biology. We used a simple cell model in the virtual microscope to rapidly prototype stimulation strategies in silico. This approach naturally extends to physically realistic models, where simulation and experiment form a closed loop: the model serves as a testbed for perturbation strategies, while experimental data parametrize and validate it. Currently, our system follows preprogrammed rules defined by the user in advance. With a sufficiently accurate model/extended control logic, the system could move beyond fixed rules and dynamically adjust experimental parameters during acquisition through active learning [105, 106], as has been demonstrated in chemistry and material sciences [107, 108]. In biology, this could translate to experiments that automatically identify rare cellular responses or even actively induce them through targeted stimulation, adjusting parameters in real time to systematically probe the underlying mechanisms. A complementary avenue is LLM-based microscope control. Modern microscopes are, in essence, imaging robots, and LLMs are already being used in robotics for natural language interfaces [109], task planning [110], and decision making [111]. In the tissue patterning experiment, we used an LLM to translate high-level strategy changes into code, but a human still evaluated the outcome; the next step is to close this loop entirely, where an LLM agent autonomously proposes, tests, and refines stimulation strategies in the (virtual) microscope, analogous to recent work in chemistry [112], with the researchers defining a high-level objective and letting agents autonomously explore the experimental space to achieve it.

## Conclusion

We demonstrated applications ranging from subcellular probing of Rho GTPase signaling at specific cytoskeletal structure and artificial recreation of traveling RhoA activity waves, to feedback-controlled assembly of cells into tissues of defined geometries and data-driven control of single-cell ERK activity in epithelial monolayers. A virtual microscope environment accelerates development by enabling *in silico* testing of feedback strategies before committing to real experiments. Looking forward, integration of natural language interfaces could broaden access to feedback-controlled microscopy, while active learning approaches could evolve this platform from rule-based automation toward autonomous experimental design.

## Code availability

The FARO software is open source (MIT license) and hosted on GitHub: https://github.com/pertzlab/FARO. Note: The pipeline was developed over the course of several years, and some experiments (Fig. 3, Fig. 4, Fig. 5 H-K) were acquired using earlier iterations of the software, including versions that used *pycro-manager* [113] rather than the current pymmcore-plus backend. The core acquisition and analysis logic remained consistent throughout development, and all experiments presented here can be reproduced with the publicly released version of the pipeline.

## ACKNOWLEDGEMENTS

This work has been supported by the Chan Zuckerberg Initiative (CZI) grant NP2-0000000095 to LH and OP, grant by the Digitalisierungskommission (Digi-K) of the University of Bern to LH, Uniscientia fellowship 187-2021 to OP, SNF Sinergia grant CRSII5_183550 to OP, Novartis Foundation grant #20C219 to OP, Faculty of Sciences of the University of Bern grant to OP, and Schweizerischer Nationalfonds (SNF) grant 310030_185376 to OP. Microscopy experiments were performed on equipment supported by the Microscopy Imaging Center (MIC), University of Bern, Switzerland.

## AUTHOR CONTRIBUTIONS

LH, AL, BG, DB and TL developed software. LH, LD and MK performed experiments. CZ established optoTrkA cell line. LH analyzed data and created the figures. LH and OP wrote the paper and acquired funding. AF, MD and OP supervised the work.

---

**Fig. S1.**
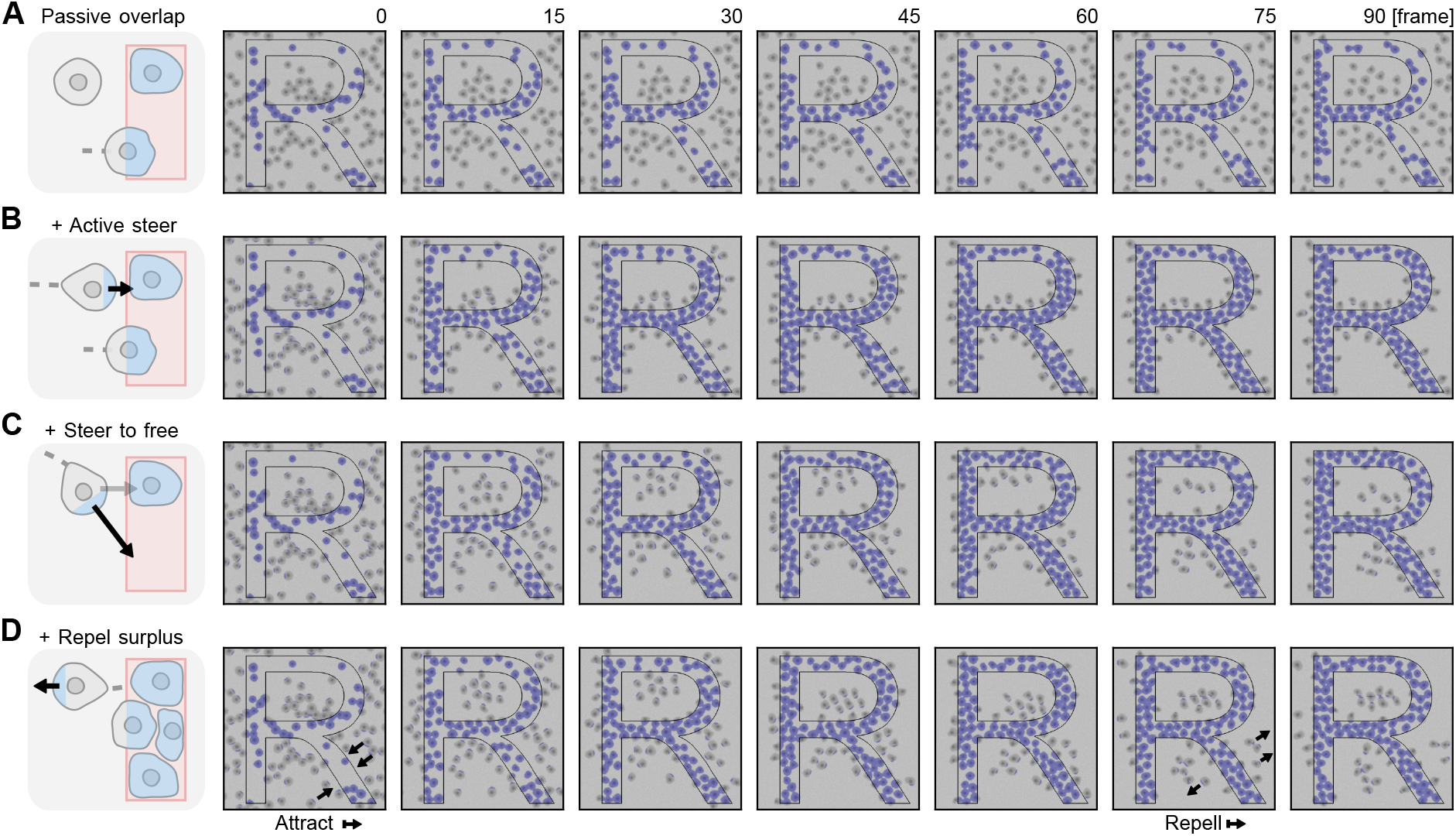
*In silico* development of stimulation strategies for tissue patterning. Stimulation strategies are evaluated and iteratively improved in a simulation before deploying the final strategy on the real microscope. Simulated cells are guided into the letter “R”. Each row shows timelapse snapshots (frames 0-90) with the stimulation mask overlaid in blue and the target outline in black. Subfigures show four strategies of increasing complexity: **A**: Passive overlap: only cells already within the target are illuminated; no active guidance. **B**: Active steer: cells outside the target are actively steered toward the reference mask, while overlapping cells are anchored in place. **C**: Steer to free: cells are directed toward *free* regions of the target, defined as the target area minus the masks of already overlapping cells expanded by a fixed radius. If no free area remains, the expansion radius is iteratively reduced until free area becomes available. **D**: Repel surplus: once an occupancy threshold is reached, cells not overlapping with the target are permanently steered away from the target, clearing the surroundings to sharpen the pattern boundary. The four strategies shown here represent illustrative steps toward the final algorithm; during actual development, many more variants were explored. After converging on the final strategy, these intermediate steps were “reconstructed” to highlight the key components of the stimulation algorithm. Associated to Fig. 2. Associated movie: **Movie S1**.

## Materials and Methods

### Cell culture

**REF52** and **NIH3T3** were maintained in DMEM high-glucose supplemented with 10% fetal bovine serum, 4 mM L-glutamine, and 100 U/ml penicillin/streptomycin. For imaging, cells were switched to FluoroBrite DMEM (Gibco) supplemented with 0.5% fetal bovine serum, 0.5% BSA, 4 mM L-glutamine, and 100 U/ml penicillin/streptomycin. For imaging, cells were seeded in 96-well 1.5H glass-bottom plates (Cellvis) coated with 10 µg/ml fibronectin (Huber lab) at a density of 1.5 x 10^3^ cells/well and incubated for 24 h. **MCF10A** human mammary cells were maintained in DMEM:F12 supplemented with 5% horse serum, 20 ng/ml recombinant human EGF (Peprotech), 10 µg/ml insulin (Sigma), 0.5 µg/ml hydrocortisone (Sigma), 200 U/ml penicillin, and 200 µg/ml streptomycin. For imaging experiments, cells were switched to a starvation medium consisting of DMEM:F12 with 0.3% BSA, 0.5 µg/ml hydrocortisone, 200 U/ml penicillin, and 200 µg/ml streptomycin. Tet-inducible optoLARG MCF10A cells were cultured in the presence of 1000 ng/ml doxycycline for 72 hours to induce transgene expression. Culture medium was replaced every 24 hours with fresh medium containing doxycycline. After 48 hours, growth medium was exchanged to starvation medium for the remaining 24 hours. For imaging, cells were seeded in 24-well 1.5H glass-bottom plates (Cellvis) coated with 5 µg/ml fibronectin (Huber lab) at 1 x 10^5^ cells/well and incubated for 48 h. All cell lines were cultured and imaged at 37°C with 5% CO_2_ and routinely tested for mycoplasma contamination by PCR and confirmed negative.

### Plasmids and cell lines

**optoLARG, 2xrGBD, Lifeact, paxillin** constructs stable cell line generation for REF52 experiments (pB3.0-optoLARG-mVenus-SspB(nano)-p2A-stargazin-mtq2-iLID-CAAX, mCherry-paxillin, dTomato-2xrGBD, Lifeact-miRFP) are described in [58].

#### optoFGFR, ERK-KTR, H2B

The optogenetic actuator/biosensor circuit for activating and reading ERK activity (optoFGFR, ERK-KTR-mRuby2, H2B-miRFP) and stable cell line generation are described in [5]. **optoTIAM** REF52 cell lines were stably transfected with CAG-optoTIAM-mVenus-iLID-nano, as described in [93].

#### optoTrkA, ERK-KTR, Lifeact, H2B

The optoTrkA plasmid was constructed using the modular cloning kit Mammalian ToolKit (MTK, Addgene kit #1000000180) [114], and additional parts from the MoClo Yeast ToolKit [115] (Addgene kit #1000000061). Transcription units (TUs) were assembled by combining MTK parts with the 678-part YTK_095 in a one-pot Golden Gate reaction with BsaI. TUs expressing PGK1-optoTrkA-mCitrine, CAG-ERKKTR-mScarlet3, CAG-H2B-miRFP670nano3, and CAG-Lifeact-miRFP670nano3 were combined into a multi-gene cassette using Esp3I and cloned into a PiggyBac expression backbone similar to MTK0_043, in which the hygromycin resistance marker was replaced by puromycin resistance. Additional basic parts (MTK3b_mScarlet3, MTK3b_miRFP670nano3, MTK3a_Lifeact) were generated by cloning the respective sequences into the domestication vector YTK001, removing all BsaI and Esp3I recognition sites by site-directed mutagenesis, and assembling them via Esp3I Golden Gate reactions. The optoTrkA construct was generated by replacing the intracellular domain of the optoFGFR1 tool [116] with a custom-synthesized intracellular domain of TrkA (Genewiz) using overhang PCR. The multi-gene PiggyBac cargo plasmid and the transposase plasmid were stably co-transfected at a 1:1 mass ratio (1 µg each) into NIH3T3 cells using jetOPTIMUS transfection reagent (Polyplus) according to the manufacturer’s protocol.

#### optoLARG, ERK-KTR, H2B

MCF10A cell lines were stably transfected with constitutively expressed CAG-ERKKTR-mRuby2-P2A-H2B-miRFP703 and Tet-inducible pTRE-sspB-mNeongreen-LARG-pTRE-Stargazin-mTurquoise2-iLID-CAG-Tet-ON. Similar to optoTrkA, the optoLARG plasmid was assembled using the modular MTK and YTK cloning libraries [114, 115]. To better accommodate the modularity of these Golden Gate based cloning kits, optoLARG was adapted from [58], and transformed into the MTK compatible parts: MTK3a_sspB-mNeongreen, MTK3a_Stargazin, MTK3b_LARG, MTK3b_mTurquoise2-iLID. To reduce cell toxicity of the construct, expression was gated through the MTK integrated doxycycline inducible promoter MTK2_001. TUs pTRE-sspB(Nano)-mNeongreen-LARG, pTRE-Stargazin-mTurquoise2-iLID, and CAG-Tet-ON-3G-transactivator were combined as described above, and integrated into the PiggyBac vector harboring the puromycin selection marker. Stable integration of the construct into MCF10A cells was done using the lonza nucleofector 2B (Lonza) using the program P-020.

Stably transfected cells were selected with puromycin (P7255, Sigma). All plasmids were verified by restriction digest and Sanger sequencing (Microsynth).

### Imaging hardware

Imaging was performed using an automated Eclipse Ti inverted fluorescence microscope (Nikon) with Nikon CFI Plan Apo Lambda 10x air (0.45 NA), CFI Plan Apo Lambda 20x air (0.75 NA), or 60x Plan Apo VC (NA 1.4) oil objective. The 1.5x magnification tube lens was engaged for experiments using the 10x objective; all other experiments used 1x. The Nikon Perfect Focus System (PFS) was used to maintain focus. Illumination was provided by a SPECTRA X LED light engine (Lumencor) with the following excitation/emission filter sets (Chroma): mCitrine: 508 nm LED, ET500/20x, 69308bs, ET535/30m; mRuby2 and dTomato: 555 nm LED, ET575/25x, 69008bs, 59022m; miRFP703: 640 nm LED, ET640/30x, 89100bs Sedat Quad, 84101m Quad. Images were acquired using an Andor Zyla 4.2 plus camera at 16-bit depth with 2x binning set for all experiments. Patterned illumination for optogenetic stimulation was projected using either an Andor Mosaic 3 or Mightex Polygon 1000 digital micromirror device (DMD). The DMD was calibrated by sequentially projecting point patterns at known DMD coordinates and detecting their positions in camera space. Detection was performed by applying Gaussian smoothing to the acquired images and locating the peak brightness coordinates. An affine transformation was then computed from three corresponding point pairs. To project patterns at desired camera-space coordinates, the inverse affine transformation was applied. Calibration accuracy was verified by projecting points at specified camera-space coordinates and visually confirming that they appeared centered at the expected locations.

### Compute hardware

Microscope computer is Intel Xeon CPU (E5-1680 v3 @ 3.20GHz), NVIDIA Quadro M4000 GPU (8GB), 64GB RAM, Windows 10 enterprise. No remote processing was used for the experiments shown, but was tested on a workstation with Intel Xeon CPU (W3-2435 @ 3.10GHz), NVIDIA RTX 4000 Ada (20GB), 128GB RAM, Windows 11 Pro. Virtual microscope simulations were run on a MacBook Pro (2021), M1 Max, 32 GB RAM, MacOS Tahoe 26.3.

### Data analysis

Nuclear markers (Fig. 1C,D, Fig. 5) acquired with 20x objective were segmented using StarDist [23] with the `2D_versatile_fluo` model (version 0.9.1). Cells with F-actin + H2B markers acquired with 10x objective (Fig. 2) were segmented using Cellpose with the `Cellpose-SAM` model (version 4.0.7) [28]. For similar experiments (not shown), we also used the `cyto3` model [27], which we recommend as a computationally lighter alternative that still leads to good segmentation results. REF52 cells imaged with 60x objective (Fig. 3, Fig. 4) were segmented using Convpaint [29], with a custom model trained on a single snapshot acquired before starting the experiment. We used a prototype version of Convpaint, using settings similar to the default model in the now public release (version 0.9.0). VGG16 pretrained on ImageNet was used as feature extractor, with the input image downscaled 2x, 16x and 32x. As the cells are relatively large and imaged at high magnification, the high downscaling factors help to capture long-range context. Random forest was used as trainable classifier implemented in `Scikit-learn` [117] with default parameters.

For all tracking tasks shown in this paper, we used `trackpy` [31] (version 0.7). Cells are tracked iteratively by initializing positions in the first frame and linking each subsequent frame’s coordinates to existing tracks, rather than reprocessing all prior frames at every timepoint. Newly assigned IDs are appended to a cumulative table across time points.

Kymographs in Fig. 3H were generated by extracting intensity profiles along a line perpendicular to the optogenetic scan at each time point, averaged over a fixed line width (50 px). To correct for photobleaching, the mean intensity across all positions was subtracted at each time point. Spatial background structure was then removed by subtracting the time-averaged intensity at each position. The resulting profiles were normalized to the mean absolute intensity. Regions outside the cell were removed using the segmentation mask. Position of the optogenetic stimulation was overlaid by tracking the centroid of the illumination mask over time.

Focal adhesion (FA) masks (Fig. 4A-E) were segmented from the normalized paxillin channel by thresholding at the 95th percentile and labeling connected components. FAs were restricted to the cell periphery, defined by eroding the cell mask by a fixed width, and filtered to exclude objects smaller than 25 or larger than 150 pixels. Each cell was divided into top and bottom halves with a line drawn perpendicular to the cell’s major axis through the centroid (determined using `Scikit-image` [118] region properties). For the FA stimulation mask, circular regions of fixed radius were placed at the centroid of each qualifying FA in the top half of the cell. For the non-FA control mask, FA regions were dilated and excluded from the cell edge to identify adhesion-free peripheral zones in the bottom half. An equal number of stimulation spots were placed at randomly selected positions within these zones, with each chosen location excluded from subsequent sampling to prevent overlap.

ERK activity quantification (Fig. 5) was performed using a multistep image processing pipeline, detailed in Dobrzyński et al. [79]. In short, mean fluorescence intensities were extracted from nuclear and cytoplasmic regions from the ERK-KTR biosensor, where the cytoplasmic region was defined as a ring surrounding each nucleus. ERK activity was quantified as the ratio of cytoplasmic to nuclear mean intensity. In Fig. 5B,F ERK activity traces were normalized to the value at the onset of stimulation for each individual cell, and short trajectories were excluded from analysis. In Fig. 5F we additionally excluded tracks exhibiting frame-to-frame changes in normalized ERK activity exceeding 40% as artifacts, and smoothed the trajectories using a rolling mean over five consecutive frames. Insets indicating the extent of ERK activity waves in Fig. 5C,G were determined by manual inspection of timelapse sequences, identifying cells that exhibited visible activation.

### Simulation virtual microscope

The simulator is wrapped as a virtual camera and SLM device implementing the pymmcore-plus UniMM-Core device API. Cells are modelled as deformable polygons (24 vertices each) whose vertex radii evolve under several concurrent forces, implemented in pygame (version 2.6.1). Each vertex radius is initialized from a randomized base radius. At each step, Brownian noise is added to the cell velocity, and the cell center is advanced with frictional damping. Vertex radii undergo stochastic membrane ruffling (low-frequency cosine perturbation), Laplacian curvature relaxation (smoothing neighbouring vertices), and radial relaxation (restoring toward the base radius). After all updates, radii are clamped and rescaled to conserve cell area. When a stimulation mask is applied, each vertex overlapping a stimulated pixel is protruded outward by a fraction of the base radius. Additionally, the cell receives a momentum impulse directed from its center toward the mean position of stimulated vertices, translating local protrusion into directed migration. A cell-cell collision is detected when the inter-center distance between two cells falls below the sum of their maximum vertex radii. Overlapping cells are separated by symmetric displacement along the center-center axis, both velocities are zeroed to prevent overlap oscillation. The simulation uses periodic (toroidal) boundary conditions. The rendered cell layer is converted to grayscale and processed through contrast reduction, Gaussian blur, additive Gaussian noise, and brightness adjustment to approximate a phase-contrast microscope image. For feedback control in the simulation, cells are segmented using a lightweight segmentation method to speed up processing: Otsu thresholding followed by a distance-transform watershed to split touching objects (input image is blurred with a gaussian filter; regions below an area threshold are discarded). In the real experiment, the segmentation method was swapped for cellpose as described above.

### Statistics

Directional bias in cell migration (Fig. 1D) was assessed by performing one-sample t-tests (scipy [119] version 1.17.0, `ttest_1samp`) against zero on the final x and y positions at 24 hours, with 95% confidence intervals computed from the standard error of the mean. Means, medians and standard errors (Fig. 1A, Fig. 4E,I, Fig. 5I) were calculated with pandas [120] (version 3.0.0).

